# Bright Cells, and Inefficient Enzymatic Activity in a Renin Hypomorph Mouse

**DOI:** 10.1101/2025.05.02.651896

**Authors:** Silvia Medrano, Lucas Ferreira de Almeida, Fernando Ruiz-Perez, Alejandro Gutierrez-Hernandez, Hiroki Yamaguchi, Daisuke Matsuoka, Manako Yamaguchi, Jason P. Smith, Thomas Wagamon, Maria Luisa S. Sequeira-Lopez, R. Ariel Gomez

## Abstract

Juxtaglomerular (JG) cells are crucial regulators of blood pressure and fluid-electrolyte homeostasis. Under normal conditions, renin secretion by JG cells is sufficient to maintain homeostasis. However, under physiological stress such as narrowing of one of the renal arteries, heart failure, dehydration, or chronic administration of renin-angiotensin system (RAS) inhibitors, additional cells along the renal arterioles are transformed to the renin phenotype to meet the demands for renin and regain homeostasis. In cases of prolonged and persistent stimulation of renin cells, concentric arteriolar hypertrophy develops. The study of renin cell identity, plasticity and function often requires the isolation of this rare cell type. Here, we report on the generation of a mouse model to label renin-expressing cells with a bright fluorescent reporter under control of the *Ren1^c^* locus for the tracking and isolation of renin cells. Kidneys from adult heterozygous (Het) *Ren1^ctdTomato/+^* mice showed tdTomato signal confined to the JG area under basal conditions, and extending along the afferent arterioles and in the intraglomerular mesangium upon treatment with captopril + low-salt diet to induce the endocrine transformation of renin cells. Unexpectedly, homozygous (Homo) *Ren1^ctdTomato/tdTomato^* mice exhibited increased tdTomato signal that extended along the afferent arterioles and into the mesangium even under normal physiological conditions, with progressive thickening of the kidney arterioles with age. Despite reduced renin immunostaining in the renal cortex, *Ren1^ctdTomato/tdTomato^* Homo mice exhibited significantly higher kidney *Ren1* mRNA and circulating renin levels when compared to Het controls. Moreover, Homo mice showed significantly lower blood pressure measured under anesthesia and angiotensin I (Ang I) plasma levels, indicating compromised renin activity. In addition, Homo mice developed interstitial fibrosis and compromised kidney function. The concentric arteriolar hypertrophy phenotype observed in these mice is identical to that described when RAS is genetically or pharmacologically inhibited, including the presence of mutations in the renin gene. Unlike mice with global deletion of renin, these animals did not require neonatal saline injections to survive and did not develop other kidney abnormalities, indicating that the bicistronic approach rendered a renin hypomorphic mouse. *Ren1^ctdTomato^* mice constitute an excellent model for the bright and strong labeling of renin-expressing cells and for the study of the mechanisms involved in the development of concentric vascular hypertrophy under RAS inhibition. In addition, this model may provide a better understanding of factors controlling renin protein folding, stability, packaging, and release.

## INTRODUCTION

Renin cells play an important role in maintaining homeostasis and are essential for survival. In adult mammals, renin cells represent a rare subset of cells – 0.01% of kidney cells. They are localized at the tip of the afferent arterioles at the entrance of the glomeruli and are therefore referred to as juxtaglomerular (JG) cells. Renin cells are responsible for the synthesis and release of renin, a critical enzyme-hormone involved in regulating blood pressure and fluid-electrolyte balance (1). Beyond a role of maintaining homeostasis, renin cells are also implicated in various physiological processes, including kidney vascular development, tissue morphogenesis, repair and regeneration, and innate immune responses (2, 3, 4, 5). During development, renin cells are precursors for cells that differentiate into smooth-muscle, mesangial, and epithelial cells, and interstitial pericytes in the kidney and show high plasticity in postnatal life in response to challenges to homeostasis (6). Under normal circumstances, secretion of renin by JG cells is sufficient to balance transient changes in blood pressure or in extracellular fluid volume. However, when there is a threat to homeostasis (such as dehydration or hypotension), renin cell descendants can reversibly revert to a renin-expressing phenotype to meet the demands for renin and restore normal physiological conditions (6). In addition, we and others have shown that conditions leading to the persistent activation of renin cells, such as experimental or spontaneous mutations of the renin-angiotensin system (RAS) genes or treatment with inhibitors of the RAS in mammals including humans, result in the development of a progressive and severe thickening of the intrarenal arterial tree characterized by the concentric hypertrophy of the kidney arterioles and arteries (7, 8, 9, 10, 11, 12, 13, 14, 15, 16).

The study of the genetic and epigenetic regulatory mechanisms underlying renin cell identity, plasticity and function often requires the identification and isolation of this rare cell type. In the past we generated a reporter mouse in which cells actively expressing renin are marked with yellow fluorescent protein (YFP) driven by a *Ren1^c^YFP* transgene (17). We have extensively used this model for *in vivo* and *in vitro* studies of renin cells (17, 18, 15). However, YFP has the disadvantage of being less bright and less photo-stable than other reporters (19, 20), hindering the ability of efficiently labeling rare cell types like renin cells. Here, we report on the generation of a mouse model that labels renin-expressing cells with the exceptionally bright fluorescent reporter tdTomato under the control of the endogenous *Ren1^c^* locus for the optimal tracking and isolation of renin cells.

## METHODS

### Generation of Ren1^c tdTomato^ mice

*Ren1^c tdTomato^* mice were generated by inserting a bicistronic *T2A-tdTomato* knock-in cassette upstream of the TGA stop codon of the mouse *Ren1^c^* gene, followed by the endogenous 3’UTR (Taconic Biosciences GmbH, Köln, Germany). To engineer the targeting vector, homology arms were generated by PCR using the BAC clone RP24-299J2 as template. In the targeting vector, a *Neo* cassette was flanked by SDA (self-deletion anchor) sites and DTA was used for negative selection. The targeting vector was subjected to restriction enzyme analysis and sequenced for confirmation purposes. The *Ren1^c^-T2A-tdTomato* targeting construct was linearized by restriction digestion with *Not*I followed by phenol/chloroform extraction and ethanol precipitation. The linearized vector was transfected into C57BL/6N ES cells according to standard electroporation procedures. The transfected ES cells were subjected to G418 selection (200μg/mL) 24 hours post electroporation. G418 resistant clones were picked and amplified in 96-well plates. The clones were subjected to DNA isolation and subsequent PCR screening for homologous recombination. Positive clones were further characterized by Southern blot analysis. Two targeted ES cell clones, 2H3 and 1A1, were injected into C57BL/6N albino embryos, which were then reimplanted into CD-1 pseudo-pregnant females. Founder animals were identified by their coat color, and their germline transmission was confirmed by breeding with C57BL/6 females and subsequent genotyping of the offspring. Heterozygous targeted mice were inter-crossed to generate homozygous targeted mice.

### Genotyping of Ren1^c tdTomato^ mice

Genotyping of the mice was conducted by PCR of DNA from tail biopsies using primers: F1: 5’-CTAGCAGATCAGAGCTTCCACAA-3’ and R1: 5’-TCACTTAGCTTCTGACCCAAAACA-3’ (PCR products size: Wildtype: 204 bp; Homozygotes: 317 bp and Heterozygotes: 317 bp/204 bp (Fig. 1A).

**Figure 1.**
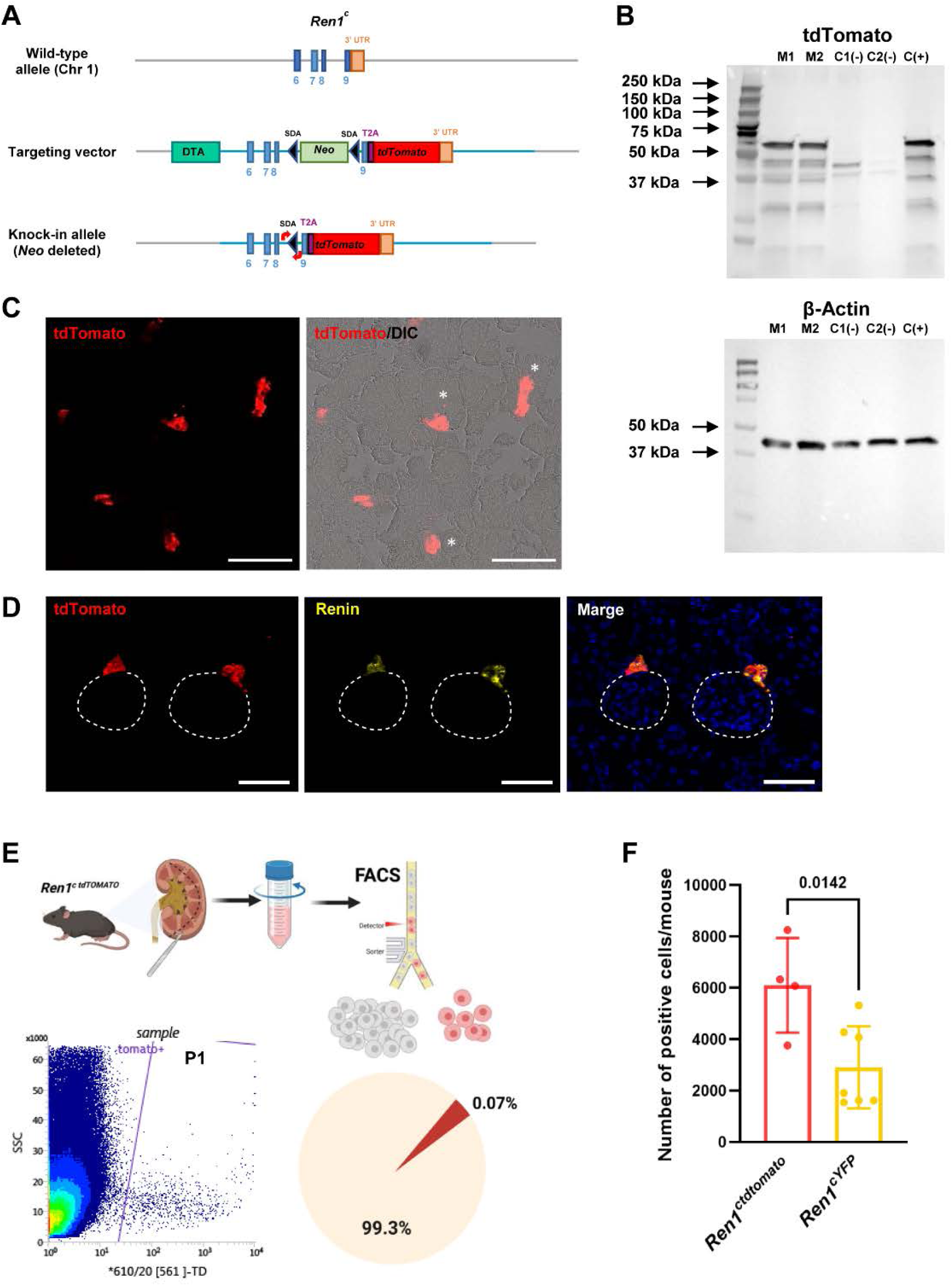
**A. Generation of *Ren1^ctdTomato^* reporter mice.** *Ren1^ctdTomato^* mice were developed by inserting a bicistronic *T2A-tdTomato* knock-in cassette upstream of the TGA stop codon of the *Ren1^c^* gene. This bicistronic construct was designed to produce Renin and tdTomato separately through a ribosomal skipping mechanism. **A**: Schematic diagram of the targeting strategy for the generation of the *Ren1^ctdTomato^* knock-in allele. ***Top Panel*.** Shown is the 3’ region of the *Ren1^c^* gene in chromosome 1 of the mouse. Exons 6-9 are indicated by numbered blue boxes. ***Middle panel*.** Map of the targeting vector. A *T2A-tdTomato* cassette was inserted immediately downstream of exon 9 of *Ren1^c^* and upstream of the endogenous 3’UTR. Positive (*Neo*) and negative (DTA) selection markers were included in the targeting vector. SDA: self-deletion anchor; DTA: diphtheria toxin A subunit; *Neo*: neomycin resistance gene. Blue lines indicate regions of homology. ***Bottom panel*.** Map of the targeted allele after *Neo* deletion. Red arrows indicate the position of the primers used for genotyping of the mice. **B. Western Blot analysis of kidney extracts from *Ren1^ctdTomato^* mice. Upper panel.** Western blot analysis for tdTomato. Kidney cortex extracts from *Ren1^ctdTomato/tdTomato^* mice (M1 and M2) were compared to kidney cortex extracts from C57BL/6 mice (C1(−) and C2(−): tdTomato negative controls) and kidney extracts from P5 *Ren1^dCretdTomato^*^/+^ mice (C+: tdTomato positive control). M1: 6 month-old *Ren1^ctdTomato/tdTomato^* female; M2: 6 month-old *Ren1^ctdTomato/tdTomato^* male; C1(−): 4-month-old C57Bl6 male; C2(−): 3-month-old C57Bl6 female; C(+): three P5 *Ren1^dCretdTomato^*^/+^ mice. Arrows indicate molecular weights. The size of the main band aligned with the predicted size of tdTomato (54.2 kDa) in both the *Ren1^ctdTomato/tdTomato^* mice (M1 and M2) and the *Ren1^dCretdTomato/+^* positive control mouse. In addition, no band of that size was present in the negative controls C1(−) and C2(−). The absence of a band of 110.6 kDa indicates that there is no formation of a Renin-tdTomato fusion protein and that the T2A system worked as expected producing two independent proteins. **Lower panel.** Western Blot analysis for the housekeeping gene β-actin was used for sample loading control. **C-F. Isolation of kidney tdTomato positive cells. C.** tdTomato expression in kidneys from *Ren1^ctdTomato/+^* mice. Shown are red fluorescence (left) and red fluorescence/DIC merged (right) images. tdTomato^+^ cells were restricted to the juxtaglomerular areas. *, glomeruli. Scale bars: 100 μm. **D.** Immunofluorescence staining confirming colocalization of tdTomato (red) and endogenous renin protein (yellow) at juxtaglomerular regions in kidney sections of *Ren1^-tdTomato+/−^* mice. Nuclei were counterstained with Hoechst (blue). Dashed circles indicate glomeruli. Scale bars: 50 µm. **E.** Fluorescence activated cell sorting (FACS) of dissociated tdTomato^+^ cells from *Ren1^ctdTomato/+^* mouse kidney cortices. ***Left panel***. Plot showing the gating strategy used to sort tdTomato+ cells. P1 demarcates cells sorted as tdTomato positive. ***Right panel***. Chart showing the percentage of tdTomato+ cells in the cell isolate. **F**. Quantification of tdTomato^+^ cells from kidneys of *Ren1^ctdTomato/+^*. The number of cells actively transcribing renin isolated from *Ren1^ctdTomato/+^* mice (6,097±1,843) was significantly higher than we previously reported for *Ren1^cYFP^* (2,902±1,594) mice (14); *p*=0.0001. Unpaired Student’s *t*-test. Data are expressed as mean ±SD.

### Animals

Both *Ren1^ctdTomato/+^* (Het) and *Ren1^ctdTomato/tdTomato^* (Homo), female and male mice were included in this study. Wild-type (Wt) *Ren1^c+/+^* mice and Ren1ctdTomato/+ (Het) were used as controls as no significant differences were observed between the two genotypes. The mice used in this study were maintained in the C57BL/6N background.

All animals were housed in the University of Virginia’s vivarium facilities equipped with controlled temperature and humidity conditions in a 12 hours dark/light cycle. All procedures were performed per the Guide for the Care and Use of Laboratory Animals published by the United States National Institutes of Health (https://grants.nih.gov/grants/olaw/guide-for-the-care-and-use-of-laboratory-animals.pdf) and approved by the University of Virginia Animal Care and Use Committee.

### Blood Pressure Measurement

Blood pressure was assessed by recording mean arterial, systolic, and diastolic values in animals under isoflurane anesthesia. Measurements were taken over a 10-minute period via a catheter inserted into the right carotid artery. The data was collected using an RX104A transducer connected to a data acquisition system and AcqKnowledge software (BIOPAC Systems, Inc., Goleta, CA).

### Blood Samples Collection and Analysis

Animals were anesthetized with an intraperitoneal injection of tribromoethanol (300 mg/kg) prior to blood collection. Blood was obtained through cardiac puncture and transferred into heparinized plasma separator tubes (BD Microtainer, Becton Dickinson, Franklin Lakes, NJ). For mice undergoing blood pressure measurements, blood was drawn from the right carotid artery while under isoflurane anesthesia. Basic metabolic panel analysis was conducted by the University of Virginia Hospital clinical laboratory.

### Plasma Renin

Plasma renin levels were measured using a mouse renin 1 ELISA kit (Ray Biotech, Norcross, GA), following the method outlined in previous studies (15).

### Glomerular Filtration Rate

The glomerular filtration rate (GFR) was measured transcutaneously using a noninvasive clearance device (NIC-Kidney Device; MediBeacon, Mannheim, Germany) and fluorescein-isothiocyanate-labelled sinistrin (FITC-S). Briefly, a small fluorescence detector was fixed on the depilated back of isoflurane-narcotized (isoflurane 4%; 0.5–1.0 L/min, O_2_) mice using a double-sided adhesive patch. After initial background fluorescent measurements, FITC-sinistrin (10 mg/100 g body weight dissolved in 0.9% NaCl) was injected intravenously and the elimination of FITC-sinistrin was measured transcutaneously in conscious mice for 60–90 minutes as previously described (21).

### Urine collection and analysis

Adult male and female 6 months-old mice were housed individually in metabolic cages designed for urine collection. After a 24 h acclimation period in metabolic cages with normal chow diet and regular water, urine samples were collected every 24 h for 2 days. Urinary albumin analysis was performed by the University of Virginia Hospital clinical laboratory, and creatinine concentration was measured using the creatinine (urinary) Colorimetric Assay kit (Cayman Chemical Co. Ltd., Ann Arbor, MI).

### Histological and Immunohistochemical Analyses

Mice were anesthetized with an intraperitoneal injection of tribromoethanol (300 mg/kg) prior to kidney removal. The kidneys were fixed overnight either in Bouin’s solution at room temperature or in 4% paraformaldehyde (PFA) at 4°C, then embedded in paraffin. Sections (5 µm) from Bouin’s-fixed, paraffin-embedded kidneys were processed for Periodic Acid-Schiff (PAS) staining (MilliporeSigma, Burlington, MA) to examine kidney morphology and picrosirius red staining to study collagen networks (Polysciences, Warrington, PA), (22).

Immunostaining was carried out on 5 µm sections of Bouin’s-fixed, paraffin-embedded kidneys using a rabbit polyclonal anti-mouse renin antibody (1:500) generated in our lab (23), or a mouse anti-α-SMA monoclonal antibody (1:10,000; MilliporeSigma), with biotinylated secondary antibodies: goat anti-rabbit IgG or horse anti-mouse IgG (1:200; Vector Laboratories, Newark, CA) for renin or α-SMA, respectively. Signal amplification was performed with the Vectastain ABC kit (Vector Laboratories) and developed using 3,3-diaminobenzidine (MilliporeSigma). Sections were counterstained with hematoxylin (MilliporeSigma), dehydrated, and mounted with Cytoseal XYL (Thermo Fisher Scientific, Waltham, MA).

tdTomato expression was observed in frozen kidney sections. Kidneys were fixed in 4% PFA for 2 h with vaccum at 4°C, washed, then incubated in 30% sucrose overnight at 4°C before being frozen in O.C.T. (Thermo Fisher Scientific, Waltham, MA). The frozen blocks were sectioned at 12 μm thickness and mounted in phosphate-buffered saline (PBS).

Immunofluorescence staining of renin was performed on 8 µm cryo-fixed sections. After blocking with 3% donkey serum, sections were incubated with anti-Renin antibody (1:5000, ab212197, Abcam, Cambridge, UK.) at 4°C overnight. After washing and blocking with 3% donkey serum, sections were incubated with Fluor 647 Donkey anti-rabbit secondary antibody (1:400, Thermo Fisher Scientific) at room temperature for 2 hours. After washing, sections were incubated with autofluorescence quenching kit (Vector Laboratories), stained for nuclei with Hoechst 33342 (Thermo Fisher Scientific) and mounted with mounting medium (Vector Laboratories).

### Western blot analysis for tdTomato

Approximately 100 mg of kidney tissue was minced with a razor blade and homogenized with a minibeater (BioSpec Products, Bartlesville, OK) using 1.0 mm zirconia beads for 2 minutes. Homogenates were lysed with RIPA buffer for 30 min on ice and sonicated for 1.5 min (30 s × 3 times, with one 30 s interval between each sonication) followed by centrifugation at 16,000×g for 20 min at 4°C. The supernatant was collected, and samples were prepared in Laemmli buffer and β-mercaptoethanol (10%) and heated at 95°C for 10 min. Samples (∼50 μg protein) were loaded onto 4–20% Tris-glycine gels (Bio-Rad) and protein samples were resolved by SDS-PAGE. Gels were transblotted onto nitrocellulose membranes for 1 h at 4°C. Thereafter, membranes were blocked with 10% milk in PBS-buffered-Tween 20 (PBST) at room temperature for 1 h and probed with goat polyclonal anti-tdTomato antibody (1:1,000; MyBiosource Inc., San Diego, CA.) or rabbit anti β-actin (1:1,500; Cell Signaling, Danvers, MA) in milk-PBST overnight at 4°C. After being washed with TBST (3 × 10 min), membranes were incubated with the corresponding secondary antibodies conjugated with horseradish peroxidase (HRP; anti-mouse or anti-rabbit) followed by PBST washes (3 × 10 min). Chemiluminescent reagent Pierce Western ECL substrate was used (ThermoFisher Scientific) to visualize protein bands using an image station. The housekeeping gene β-actin was used for sample loading control.

### Microscopy

Kidney sections were visualized using a Zeiss Imager M2 microscope equipped with AxioCam 305 color and AxioCam 506 mono cameras (Zeiss, Oberkochen, Germany).

### RNA isolation and Real-time RT-PCR analysis

RNA isolation and quantitative Real-Time PCR (qPCR) analysis were carried out on kidney cortex samples. Total RNA was extracted using TRIzol reagent (Thermo Fisher Scientific) and the RNeasy Mini Kit (Qiagen, Germantown, MD). Reverse transcription was performed with oligo(dT) primers and M-MLV Reverse Transcriptase (Promega, Madison, WI) at 42°C for 1 hour, following the manufacturer’s guidelines. qPCR was conducted using SYBR Green I (Thermo Fisher Scientific) in the CFX Connect system (Bio-Rad Laboratories, Hercules, CA). The PCR reactions were run with the following primers: *Ren1*, forward: 5′-ACAGTATCCCAACAGGAGAGACAAG-3′, reverse: 5′-GCACCCAGGACCCAGACA-3′; *Rps14*, forward: 5′-CAGGACCAAGACCCCTGGA-3′, reverse: 5′-ATCTTCATCCCAGAGCGAGC-3′. Expression of *Ren1* mRNA was normalized to *Rps14*, and fold changes in expression were determined using the ΔΔCt method, with results reported as relative expression compared to control mice.

### Isolation of tdTomato expressing cells

To measure the number of tdTomato-positive cells in *Ren1^ctdTomato^* mouse kidney cortices, we used fluorescent activated cell sorting (FACS). The kidneys were excised and decapsulated, and the cortices were dissected, minced, and transferred into a VIA Extractor™ tissue pouch (Cytiva, Marlborough, MA) containing 5 mL of enzymatic solution (0.3% collagenase A, 0.25% trypsin and 0.0021% DNase I, all from MilliporeSigma). The pouches were placed inside a VIA Extractor™ tissue disaggregator (Cytiva) and the tissue was disaggregated at 200 rpm for 12 minutes at 37°C. The tissue was centrifuged at 800×g for 4 minutes at 4°C. The cell pellet was resuspended in fresh Buffer 1 (130 mM NaCl, 5 mM KCl, 2 mM CaCl_2_, 10 mM glucose, 20 mM sucrose, 10 mM HEPES, pH 7.4), and the suspension was poured consecutively through sterile 100 μm and 40 μm nylon cell strainers (Corning Inc., Corning, NY). The flow-through was centrifuged at 800×g for 4 minutes at 4°C. Residual red blood cells were lysed with Red Blood Cell Lysis Buffer (MilliporeSigma). The cell pellet was resuspended in 1.5 mL of PBS, 1% fetal bovine serum (FBS), 1 mM ethylene diamine tetraacetic acid (EDTA), and DNase I (MilliporeSigma). The dead cells were labeled with DAPI (MilliporeSigma). Cells were analyzed and sorted for 2 hours using an Influx Cell Sorter (Becton Dickinson, Franklin Lakes, NJ) at the Flow Cytometry Core Facility of the University of Virginia. Finally, cells were collected in DMEM, 20% FBS for further analysis.

### Statistical Analysis

Data are presented as means ± SD. Statistical analysis was performed using GraphPad Prism 10 software (GraphPad Software, San Diego, CA). Comparisons between two groups were performed by two-tailed unpaired Student’s *t*-test. *P* values < 0.05 were considered statistically significant.

## RESULTS

### Renin cells are efficiently labelled in Ren1^ctdTomato^ mice

To label cells actively expressing renin, we developed a knock-in mouse in which the expression of the fluorescent reporter tdTomato is driven by the endogenous *Ren1^c^* locus. *Ren1^ctdTomato^* mice were generated by inserting a bicistronic *T2A-tdTomato* knock-in cassette upstream of the TGA stop codon of the *Ren1^c^* gene (Fig. 1A). Multicistronic constructs are commonly used to express reporter genes in mice. They consist of two or more genes linked by 2A peptide sequences, which produce separate proteins through a ribosomal skipping mechanism (24). To determine the efficacy of the bicistronic approach in our mouse model, we conducted Western Blot analysis of kidney extracts from *Ren1^ctdTomato/tdTomato^* Homo mice, *Ren1^dCretdTomato/+^* positive control (a strain that expresses tdTomato following Cre-mediated recombination in renin cells) and C57BL/6 (tdTomato negative control) mice (Fig. 1B). A 54.2 kDa band would indicate the presence of tdTomato. If the bicistronic knock-in did not work efficiently, a 116 kDa band would appear indicating a Renin-tdTomato fusion protein. Our results show that the size of the main band aligned with the predicted size of tdTomato (54.2 kDa) in both the *Ren1^ctdTomato/tdTomato^* mice (M1 and M2) and the *Ren1^dCretdTomato/+^* control mouse. In addition, no band of that size was present in the negative controls, confirming the specificity of the tdTomato antibody. Importantly, we did not observe a 116 kDa band in *Ren1^ctdTomato/tdTomato^* mice, indicating that there was no Renin-tdTomato fusion protein generated and that the T2A system worked as expected producing two independent proteins.

Next, we sought to determine if *Ren1^ctdTomato^* mice can be used to effectively isolate cells actively transcribing renin. Figure 1C shows images of a kidney frozen section from a *Ren1^ctdTomato/+^* mouse. tdTomato signals can be clearly and exclusively seen in the JG area of the kidney. To measure the number of tdTomato-positive cells in the kidney, we isolated cells from kidney cortices of *Ren1^ctdTomato/+^* mice using FACS (Fig. 1D). The percentage of tdTomato^+^ cells in the cell isolate was (0.063±0.017%). The number of cells actively transcribing renin per mouse isolated from *Ren1^ctdTomato/+^* mice was significantly higher than we previously reported for *Ren1^cYFP^* mice (14), 6,097±1,843 vs 2,902±1,594 (*p*=0.0142) (Fig. 1E).

These results demonstrate that tdTomato allows clear identification and quantification of cells with an active renin promoter, highlighting the advantage of using a brighter, and more photostable reporter for labeling renin-expressing cells.

### Kidneys of Ren1^ctdTomato/+^ heterozygous mice exhibit a normal distribution of renin-expressing cells

We first investigated the pattern of tdTomato expression in frozen kidneys from adult heterozygous (Het) *Ren1^ctdTomato/+^* mice. We found tdTomato signal confined to the JG area under basal conditions (Fig. 2A). In addition, tdTomato signal extended along the afferent arterioles and in the intraglomerular mesangium upon treatment with captopril + low-salt diet to induce the endocrine transformation of renin cells (Fig. 2B). This pattern of tdTomato expression closely resembled the expression of endogenous renin revealed by immunostaining in both the basal state and under homeostatic stress (Fig. 2C and D).

**Figure 2.**
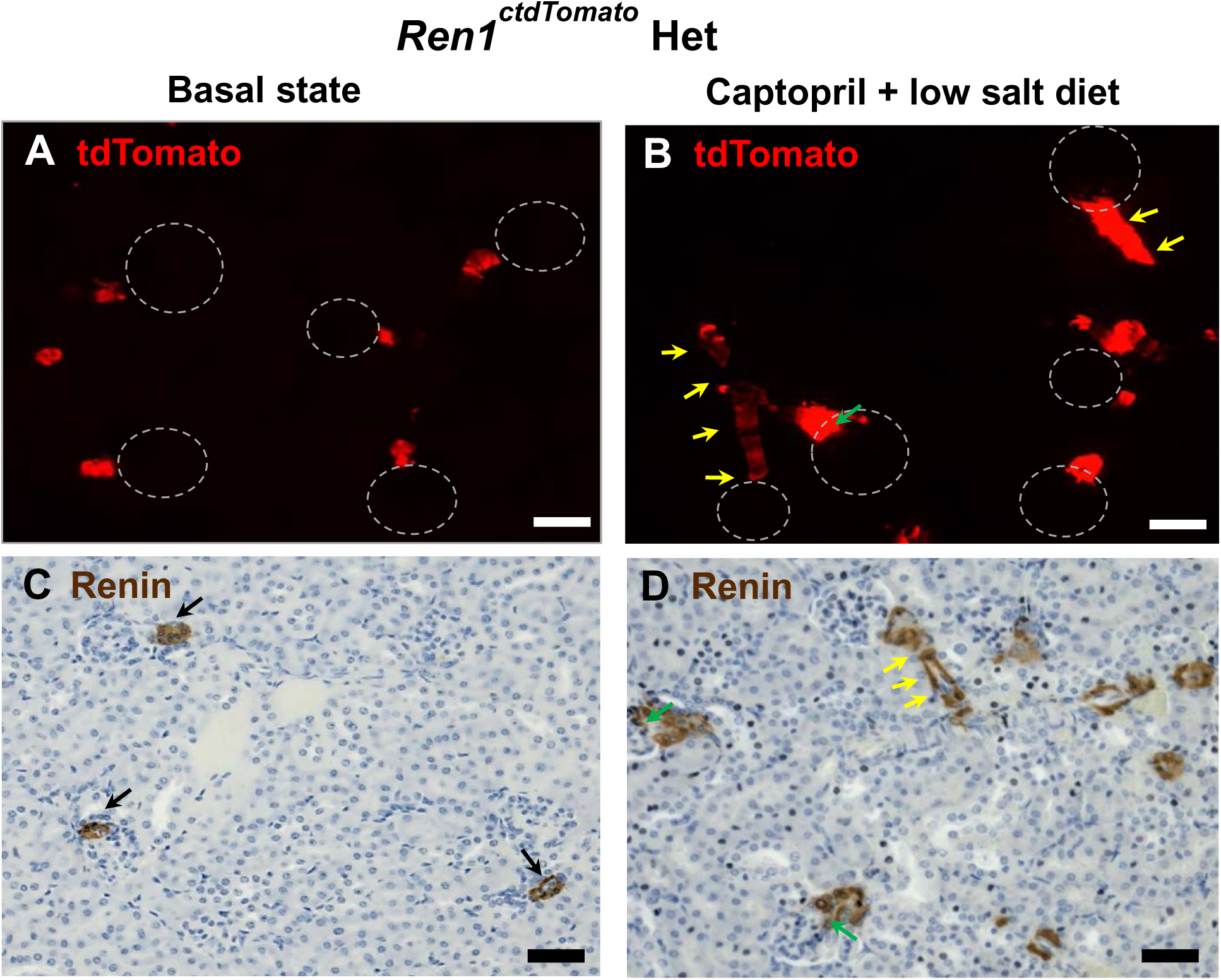
tdTomato and Renin expression in adult *Ren1^ctdtomato^* heterozygous mouse kidneys. **A-B.** tdTomato expression in kidney frozen sections. **A.** Under normal physiological conditions, tdTomato signal was confined to JG areas. White dashed circles indicate glomeruli. Bar: 50 µm. **B.** In mice treated with 0.5 mg/mL of captopril in the drinking water and fed a low-NaCl (0.1%) diet for 8 days to induce the endocrine transformation of renin cells, tdTomato expression extended along the afferent arterioles (yellow arrows) and the intraglomerular mesangium (green arrows). White dashed circles indicate glomeruli. Bar: 50 µm. **C-D.** Renin immunostaining of kidney sections from the same animals shown in A and B. **C.** Results show the normal renin distribution shown in brown (black arrows) at the entrance of the glomeruli and in the arterioles under basal conditions. **D.** In response to Captopril + low Na^+^ diet treatment, kidneys showed renin along the arterioles (yellow arrows) and in the mesangium (green arrows). Bar: 50 µm. These results indicate that, in *Ren1^ctdtomato^* heterozygous, the pattern of tdTomato expression is identical to that of endogenous renin under both basal conditions and homeostatic stress. Mice: 60 days-old females.

These results indicate that *Ren1^ctdTomato/+^* mice constitute a robust model for the identification and localization of actively expressing renin cells.

### Ren1^ctdTomato^ homozygous mice exhibit low levels of renin in the kidney and progressive thickening of the kidney arteries and arterioles

Next, we analyzed *Ren1^ctdTomato/tdTomato^* Homo mice at 2 months of age Unexpectedly, unlike *Ren1^ctdTomato/+^* Het animals, mice with two copies of the knock-in *T2A-tdTomato* construct exhibited increased tdTomato signal that extended in the afferent arterioles and the mesangium under normal physiological conditions (Fig. 3A). In addition, the vasculature exhibited abnormalities, characterized by apparent thickening of the vessel walls. We observed similar atypical tdTomato expression and vessel structure in Homo mice subjected to captopril and low-salt diet for a week (Fig. 3B). Despite the increased tdTomato signal in *Ren1^ctdTomato/tdTomato^* Homo mouse kidneys, we observed a severe reduction in renin expression by immunostaining under both normal physiological conditions and homeostatic stress (Fig. 3C and D).

**Figure 3.**
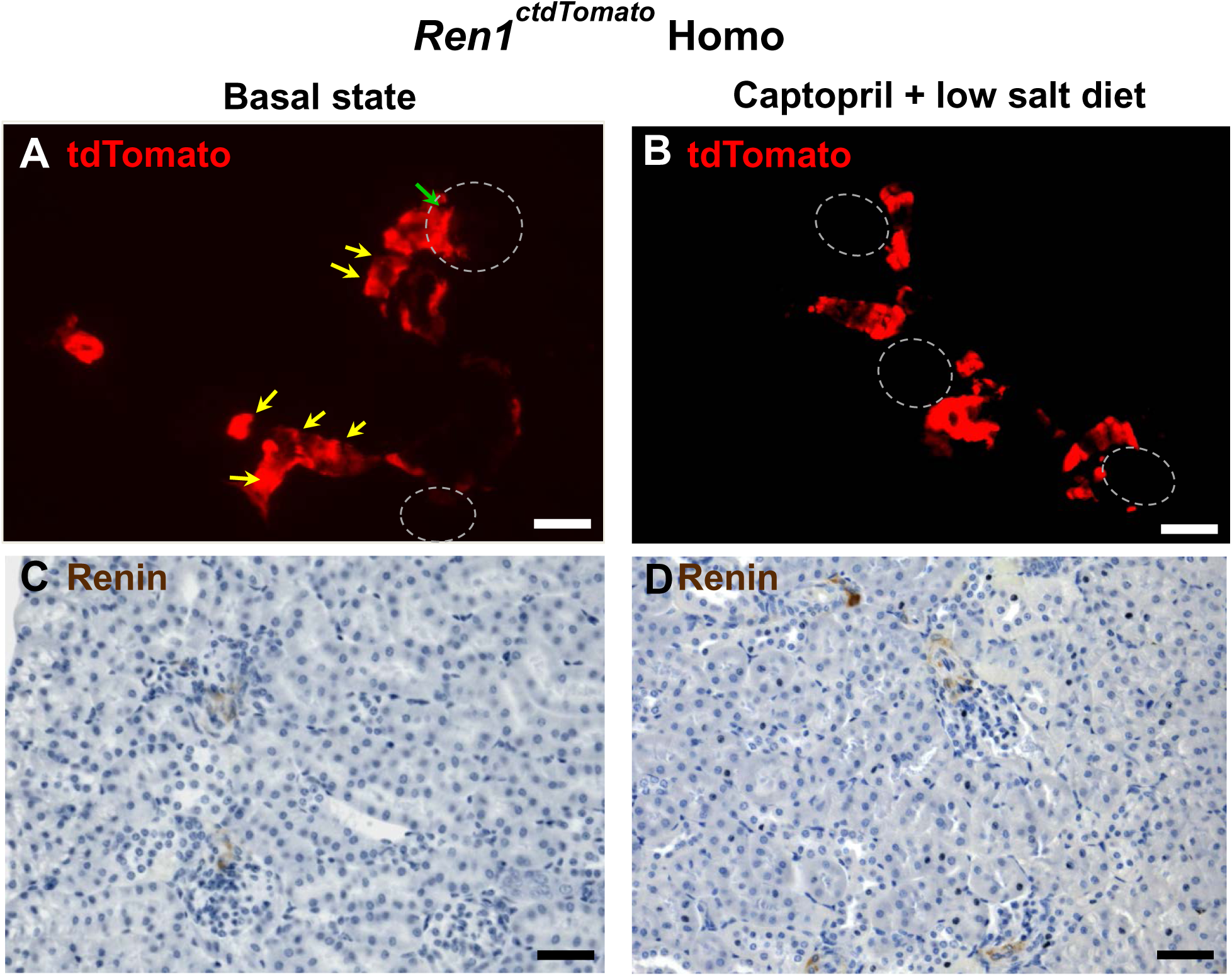
tdTomato and Renin expression in adult *Ren1^ctdtomato^* homozygous mouse kidneys. **A-B.** tdTomato expression in kidney frozen sections. **A.** Unexpectedly, under normal physiological conditions, mice with two copies of the knock-in *T2A-tdTomato* construct exhibited increased tdTomato signal that extended in the afferent arterioles (yellow arrows) and the mesangium (green arrows). In addition, the vasculature was abnormal with apparent thickening of the vessel walls. White dashed circles indicate glomeruli. Bar: 50 µm. **B.** Similar atypical tdTomato expression and vessel morphology was observed in *Ren1^ctdtomato/tdTomato^* Homo mice subjected to captopril and low-salt diet. Glomeruli are indicated by white dashed circles. Bar: 50 µm. **C-D.** Renin immunostaining of kidney sections from the same animals shown in A and B. **C.** Results show that, despite the increase in tdTomato signal in these animals, there was a severe reduction in Renin expression by immunostaining under both normal physiological conditions and homeostatic stress. Bars: 50 µm. Mice: 60 days-old females.

To attest in detail the morphology of the vasculature of *Ren1^ctdTomato/tdTomato^* Homo kidneys, we conducted immunostaining for α-smooth muscle actin (ACTA2). At 2.5 months of age, kidneys from Homo mice showed thicker arteriolar walls than Het counterparts accompanied by reduced vessel lumen (Fig. 4A, B). In addition, this vascular phenotype appeared to progress over time. At 4-5 months of age, *Ren1^ctdTomato/tdTomato^* Homo mice exhibited further hypertrophy of the arterioles and arteries with significant narrowing of the vascular lumen, and positive ACTA2 intraglomerular and peritubular staining (Fig. 4C-F). This phenotype resembles the one observed with spontaneous mutations of RAS genes or long-term treatment with RAS inhibitors.

**Figure 4.**
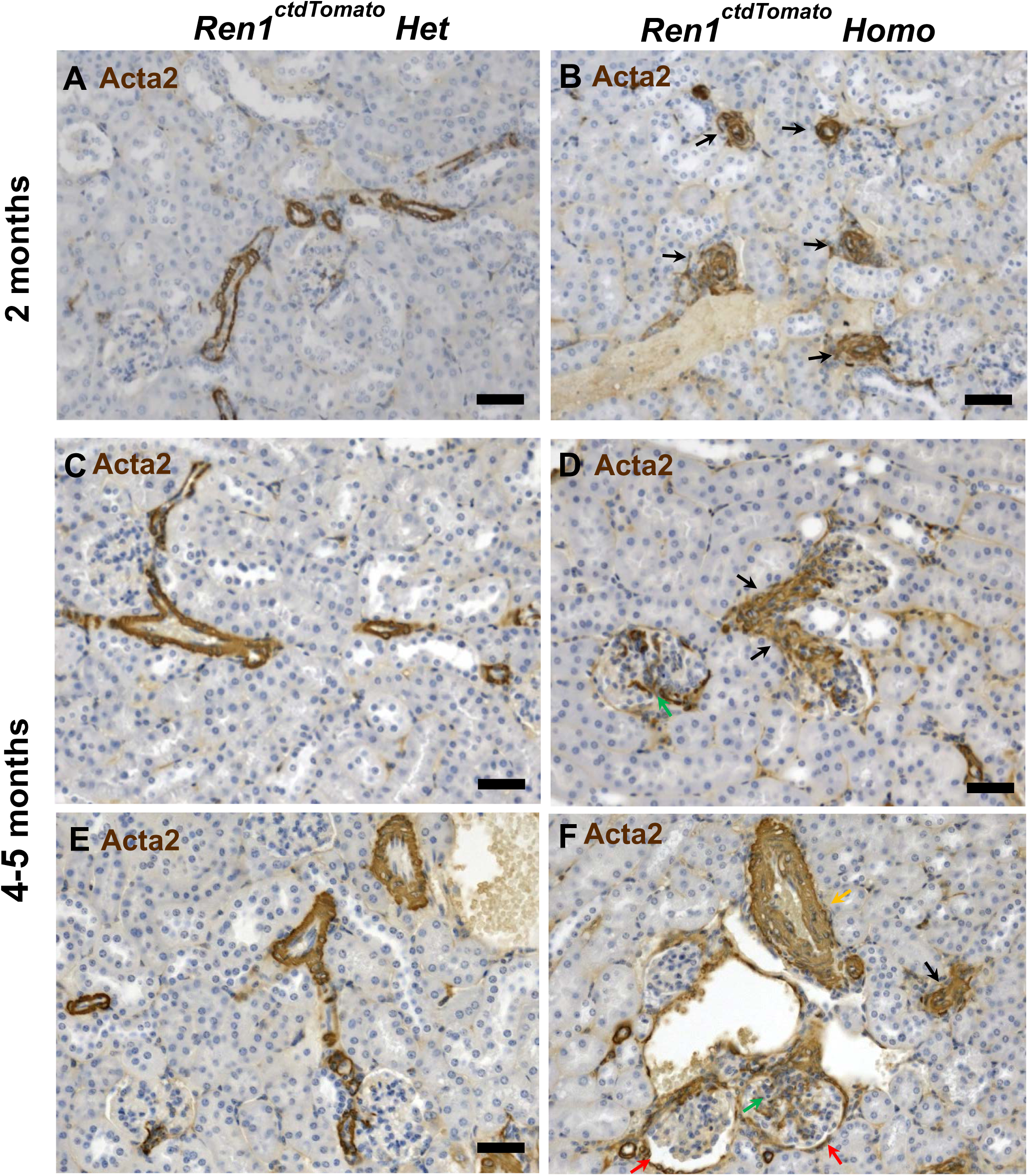
*Ren1^ctdtomato^* homozygous mice exhibit progressive thickening of arterioles and arteries. Shown are images of immunostaining for Acta2 (labeled in brown) of kidney sections from *Ren1^ctdTomato^* heterozygous (**A, C, E**) and homozygous (**B, D, F**) mice. **A, B.** At 2 months of age, kidneys from *Ren1^ctdTomato/tdTomato^* Homo mice showed thicker arteriolar walls (black arrows) than Het counterparts. Bars: 50 µm. Mice: 60 days-old males. **C, D, E, F.** At 4-5 months of age, *Ren1^ctdTomato^* Homo mice exhibited further hypertrophy of the arterioles (black arrows) and arteries (yellow arrows) with significant narrowing of the vascular lumen, in addition to positive ACTA2 intraglomerular (green arrows) and peritubular staining (red arrows). Bars: 50 µm. Mice: 150 days-old male (**C**), 167 days-old male (**D**), 150 days-old female (**E**), 167 days-old female (**F**). This phenotype resembles the one observed with spontaneous mutations of renin-angiotensin system (RAS) genes or long-term treatment with RAS inhibitors.

**Figure 5.**
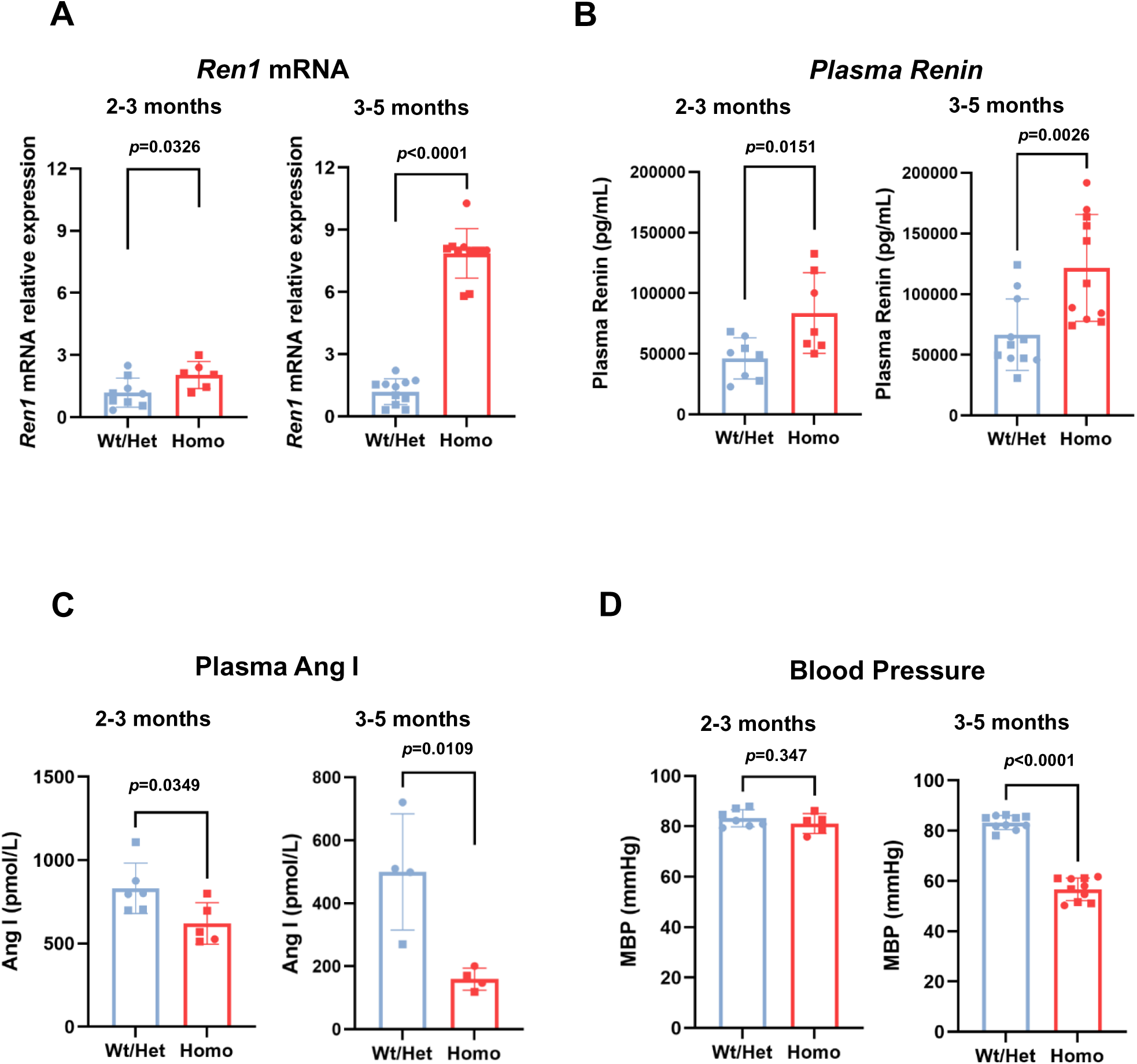
*Ren1^ctdTomato^* homozygous mice exhibit high levels of *Ren1* mRNA and circulating Renin, reduced levels of Ang I, and low blood pressure. **A.** *Ren1* mRNA levels in kidney cortices. We observed a significant increase in renin mRNA levels in *Ren1^ctdTomato/tdTomato^* Homo mice under normal physiological conditions compared to Wt/Het controls that progressed with age. Age: 2-3 months: Wt/Het, n=9; Homo, n=6, *p*=0.0326. Age: 3-5 months: Wt/Het, n=11; Homo, n=11, *p*<0.0001. Unpaired Student’s *t*-test. Squares: males; circles: females. **B.** Plasma renin. Circulating renin levels followed a similar trend despite the marked reduced levels of Renin expression at the protein level in the kidney. Age: 2-3 months: Wt/Het, n=8; Homo, n=7, *p*=0.0151. Age: 3-5 months: Wt/Het, n=11; Homo, n=11, *p*=0.0026. Unpaired Student’s *t*-test. Squares: males; circles: females. **C**. Plasma Ang I. Circulating Ang I levels were significantly lower in Homo mice compared to Wt/Het counterparts. Age: 2-3 months: Wt/Het, n=6; Homo, n=6, *p*<0.0349. Age: 3-5 months: Wt/Het, n=11; Homo, n=11, *p*<0.0109. Unpaired Student’s *t*-test. Squares: males; circles: females. **D**. Arterial blood pressure. *Ren1^ctdTomato/tdTomato^* Homo mice exhibited a significant decrease in arterial blood pressure compared to controls at 3-5 months of age. Age: 2-3 months: Wt/Het, n=7; Homo, n=5, *p*=0.347. Age: 3-5 months: Wt/Het, n=10; Homo, n=10, *p*=0.0001. Unpaired Student’s *t*-test. Squares: males; circles: females. MBP, mean blood pressure. Data are expressed as mean ± SD for all measures.

### *Ren1^ctdTomato/tdTomato^* homozygous mice exhibit high levels of kidney *Ren1* mRNA and circulating Renin, reduced renin activity and low blood pressure

Next, we examined the expression of renin at the mRNA level in the kidney cortex of *Ren1^ctdTomato^* mice by qPCR. We observed that, despite reduced renin protein in the kidney, *Ren1^ctdTomato/tdTomato^* Homo mice exhibited significantly higher *Ren1* mRNA compared to Wt/Het controls at both 2-3 months (2.04 ± 0.65 vs 1.12 ± 0.70, *p*=0.0326) and 3-5 months (7.86 ± 1.19 vs 1.20 ± 0.62, *p*<0.0001) of age, suggesting that the animals are attempting to increase renin production. In addition, we found that the levels of circulating renin followed a similar trend at both 2-3 months: (83,645 ± 33,325 vs 46,613 ± 16,994, *p*=0.0151) and 3-5 months: (121,737 ± 44,005 vs 66,731 ± 29,411, *p*=0.0026). On the other hand, circulating Ang I levels were significantly lower in Homo mice compared to Wt/Het counterparts at 2-3 months: (830.4 ± 151.5 vs 620.1 ± 123.9), *p*=0.0349) and 3-5 months: (500.0 ± 184.1 vs 159.3 ± 34.8, *p*=0.0109), indicating compromised renin activity. In line with the reduced levels of Ang I in plasma, we found that *Ren1^ctdTomato/tdTomato^* Homo mice exhibited a significant decrease in arterial blood pressure compared to controls at 3-5 months of age: (83.27 ± 2.77 vs 56.72 ± 4.53, *p*<0.0001). At 2-3 months of age, blood pressure was not significantly changed (83.20 ± 3.37 vs 81.12 ± 3.90, *p*=0.3470).

These results indicate that *Ren1^ctdTomato/tdTomato^* Homo mice have compromised renin activity that results in elevated levels of mRNA in the kidney and of circulating renin to overcome renin deficiency. At the same time, reduced renin activity results in a significant reduction of blood pressure.

### Ren1^ctdTomato/tdTomato^ homozygous exhibit interstitial fibrosis and compromised kidney function

We next examined the morphological structure of the kidney in the *Ren1^ctdTomato^* model. PAS staining of full kidney sections revealed preserved overall architecture, with no visible abnormalities in the cortex, outer medulla, inner medulla or papilla in *Ren1^ctdtomato^* Homo mice (Fig. 6A). However, Picrosirius red staining revealed signs of vascular remodeling, including increased collagen deposition in the vessel walls, intraglomerular and periglomerular areas, and the interstitium (Fig. 6B, left panels). Immunohistochemical analysis for Acta2 of the same mice demonstrated significant arteriolar wall thickening and vascular remodeling (Fig. 6B, middle panels). Similarly, renin immunostaining showed a marked reduction in renin expression in *Ren1^ctdtomato^*^/tdTomato^ Homo animals compared to Wt/Het counterparts (Fig. 6B, right panels). Functional evaluation of *Ren1^ctdtomato^* Homo mice revealed a significant reduction in the transdermal glomerular filtration rate (tGFR, Fig. 6C), accompanied by significantly prolonged FITC-sinistrin half-life (Fig.6D), and elevated blood urea nitrogen (BUN, Fig. 6E) compared to WT/Het controls. These results are consistent with compromised renal function in *Ren1^ctdtomato^* Homo mice. Despite these functional changes, the urine albumin-to-creatinine ratio (UACR) remained unchanged between the groups, indicating preserved glomerular barrier function (Fig. 6F).

**Figure 6.**
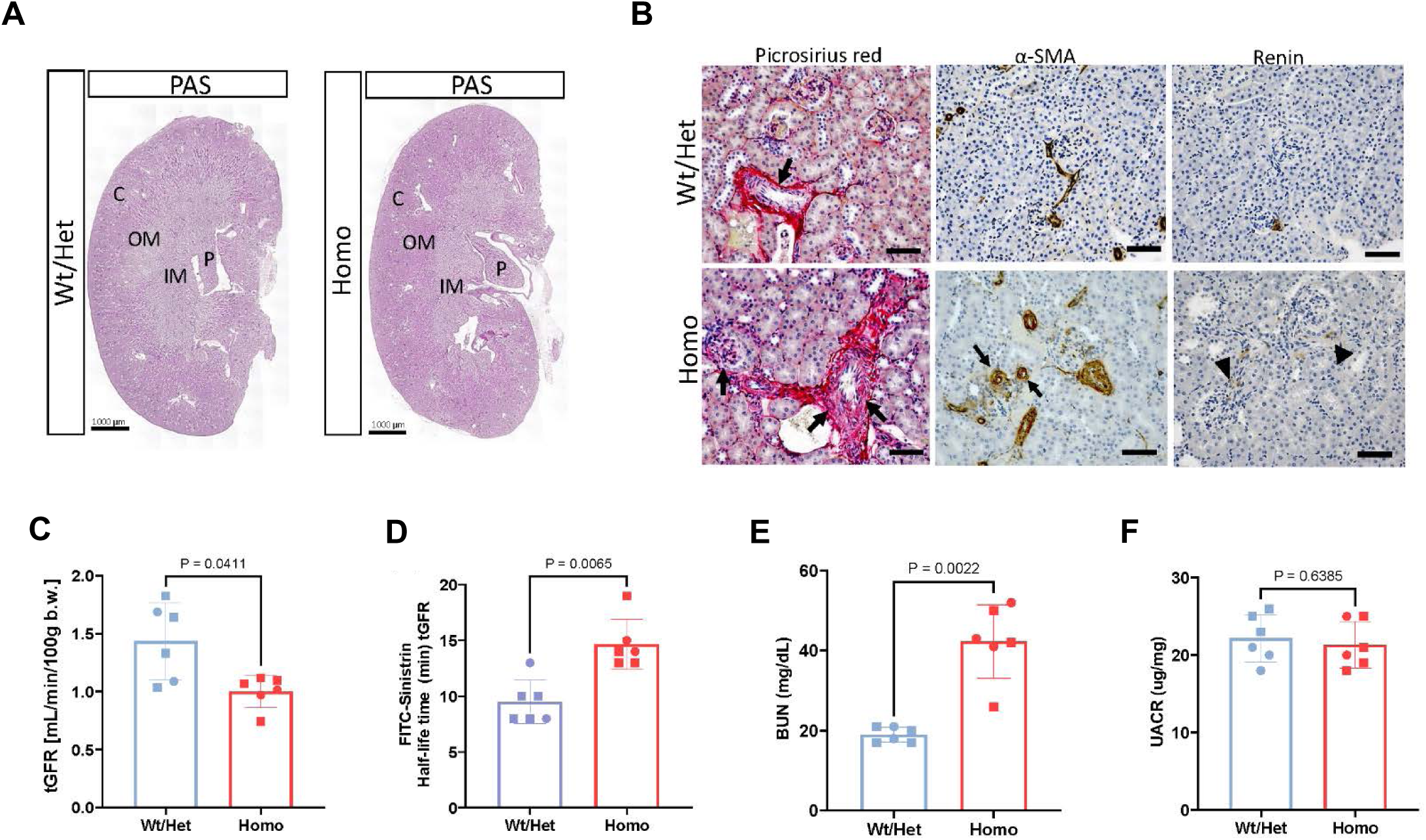
*Ren1^ctdTomato^* homozygous mice exhibit compromised kidney function. **A.** Representative light microscopy images of periodic acid-Schiff (PAS) stained kidney sections from 6-month-old mice reveal preserved overall renal architecture with no overt morphological differences between *Ren1^ctdTomato^* Homo and Wt/Het mice. C, Renal cortex; OM, Outer medulla; IM, Inner medulla; P, Papilla. Bars: 1 mm **B. Left panels.** Picrosirius red staining revealed histological changes consistent with vascular remodeling and accumulation of collagen in vessel walls, intraglomerular and periglomerular areas, and the interstitium (black arrows). Bars: 50 µm. **Middle panels.** Immunohistochemical staining for Acta2 showed significant arteriolar wall thickening and perivascular signal accumulation. Bars: 50 µm. **Right panels**. Immunohistochemistry for renin revealed a marked decrease in renin expression in the JG area of Homo mice compared to Wt/Het counterparts (arrowheads). Bars: 50 µm. **C-E.** *Ren1^ctdTomato^* Homo mice exhibited impaired renal function demonstrated by significant reduction in (C) transdermal glomerular filtration rate (tGFR), (D) FITC-sinistrin half-life, and (E) blood urea nitrogen (BUN). **F.** Urine albumin-to-creatinine ratio (UACR) did not differ significantly between the groups. Data are expressed as mean ± SD, n= 6 per group.

## DISCUSSION

In this study, we describe a novel reporter mouse that labels renin-expressing cells with a bright fluorescent reporter under the control of the *Ren1^c^* locus, enabling the tracking and isolation of renin cells. This model consists of a knock-in *T2A-tdTomato* cassette inserted immediately upstream of the TGA stop codon of the mouse *Ren1*^c^ gene. The *Ren1^ctdTomato^* model has significant advantages over existing renin cell reporter lines, including our *Ren^cYFP^* model (17). In particular, tdTomato, a tandem dimer derived from the original red fluorescent protein DsRed, is the brightest dimer fluorescent protein reporter available, with more than twice the brightness of eYFP (19). In addition, tdTomato is highly photostable and less sensitive to external factors than other reporters (20). Previous studies have shown that YFPs, for example, show excessive pH sensitivity, chloride interference, poor photostability, and poor expression at 37 °C (25). Indeed, our results showed that twice as many renin-expressing cells can be isolated from the kidney of *Ren1^ctdTomato^* mice compared to the *Ren^cYFP^* model. This represents a substantial improvement in the study of extremely rare JG cells, which constitute only about 0.01% of the kidney’s cellular mass. Although different dissociation methods were used—VIA Extractor in this study and manual dissociation previously— both approaches proved to be highly efficient in disaggregating the tissue. The higher number of renin cells observed is therefore likely due to the increased sensitivity of the tdTomato reporter.

In addition to the above-mentioned advantages, a knock-in experimental approach avoids potential problems associated with transgenic lines that are prone to spurious and/or ectopic gene expression (26). Since the *T2A-tdTomato* cassette was inserted upstream of the TGA of the endogenous *Ren1^c^* gene, tdTomato expression is expected to be regulated by the same elements that control renin expression. Indeed, our results show that the red fluorescence signal in the kidneys from heterozygous *Ren1^ctdTomato/+^* mice clearly resembles the pattern of endogenous renin protein expression in the adult under basal conditions and when homeostasis was compromised.

Overall, the *Ren1^ctdTomato^* mouse is a highly specific and sensitive model for studying renin-expressing cells and represents a superior tool for animal studies requiring a bright and stable reporter signal, particularly for *in vivo* imaging and high-resolution 3D tissue morphology analysis.

An unexpected finding in our study is that mice homozygous for the *T2A-tdTomato* cassette developed progressive concentric arterial and arteriolar hypertrophy (CAAH), resulting in stenosis of the vessels. This phenotype is reminiscent of the vascular pathology we and others observed in mammals, including humans, with experimental or spontaneous mutations of any of the RAS genes or long-term treatment with inhibitors of the RAS (7–16). All these manipulations result in the overactivation of renin cells to increase renin release and counterbalance the deficit in RAS function and re-establish homeostasis. We have reported that renin cells *per se* contribute directly to the vessel thickening (15, 27), not by increased proliferation (28) but rather by their transformation from an endocrine to an embryonic invasive matrix-secretory phenotype characterized by the production of osteopontin, biglycan, matrix gla protein, angiogenic molecules, growth factors, and cytokines, accompanied by inward accumulation of transformed smooth muscle cells (15). Nevertheless, the underlying mechanisms leading to this vascular pathology remain unknown and the *Ren1^ctdTomato^* mouse model opens a new avenue for the study of this disease.

The increased tdTomato signal in *Ren1^ctdTomato^* Homo mouse kidneys indicates that the renin transcriptional machinery is activated in these mice. This was corroborated by the presence of significantly higher levels *Ren1* mRNA in the kidney. Despite the renin transcriptional activation, we observed a severe reduction in Ren1 protein in the kidney under both normal physiological conditions and homeostatic stress. On the contrary, the levels of renin in the circulation were significantly higher in *Ren1^ctdTomato^* Homo mice, accompanied by a significant reduction in Ang I. The latter suggests that these mice have impaired renin activity. It is possible that the increased levels of *Ren1* mRNA represent an attempt by renin-producing cells to counterbalance reduced renin activity by producing more renin. The reduced renin levels in the kidney, accompanied by increased circulating renin, may indicate enhanced release in response to diminished renin activity.

It has been reported that with the T2A system, complete separation of the two proteins is not always achieved, and some protein fusions can still occur (29, 30). Therefore, one explanation for the apparent loss of renin activity is that the bicistronic mechanism used for the generation of this model is not working efficiently to produce separate renin and tdTomato proteins, generating instead a fusion protein with compromised activity. Our Western blot results, however, argue against this conclusion. We did not detect the presence of a Renin-tdTomato fusion protein, indicating that in our model the *Ren1c-T2A-tdTomato* transcript is efficiently translated into Renin and tdTomato as separate proteins by skipping the glycine-proline peptide bond formation at the C-terminus of the 2A peptide. Another consequence of the T2A bicistronic approach is that because of the ribosomal skipping mechanism, 20 amino acids (GSGEGRGSLLTCGDVEEN) of the T2A peptide are added to the C terminus of the upstream protein, Renin, while one proline residue is added to the N terminus of the downstream protein, tdTomato. It is reasonable to conclude that the extra amino acids added to the C terminus of Renin may produce a conformational change leading to abnormal folding, compromised enzymatic activity, and/or reduced stability of the protein. Future studies involving protein 3D structure prediction tools using the Renin-T2A sequence will shed light on the mechanisms involved in the generation of the abnormal renin protein, and provide a better understanding of factors controlling renin protein folding, stability, packaging, and release.

Our model, characterized by impaired renin activity in *Ren1^ctdTomato/tdTomato^* Homo mice, exhibited significant vascular remodeling, including concentric arteriolar hypertrophy and a marked reduction in renin expression within the JG region. Despite these vascular abnormalities, histological analysis revealed preserved renal architecture across all regions, including the cortex, outer medulla, inner medulla, and papilla. Functionally, *Ren1^ctdTomato/tdTomato^* Homo mice demonstrated impaired renal function, as evidenced by reduced glomerular filtration rate and elevated BUN levels, and were hypotensive, consistent with diminished RAS activity. Previous studies have shown that angiotensin II (Ang II) signaling through the angiotensin II type 1 (AT1) receptor is essential for postnatal development of the renal microvasculature (31). Disruption of this pathway during nephrogenesis has been linked to severe medullary atrophy and hydronephrosis, as well as defective development of the peristaltic apparatus and pressure-induced epithelial injury (32). The preservation of medullary architecture in our model suggests that RAS activity during kidney development was not sufficiently impaired to disrupt medullary formation. However, in adulthood, reduced systemic blood pressure appears to trigger chronic recruitment of renin-lineage progenitor cells in an attempt to restore homeostasis. This persistent activation drives pathological vascular remodeling, ultimately resulting in the CAAH observed in our model. Furthermore, picrosirius red staining revealed increased collagen deposition not only in the vasculature but also within glomeruli. This finding aligns with studies suggesting that renin-lineage mesangial cells can undergo phenotypic switching under chronic stimulation, re-expressing renin and acquiring a secretory profile that contributes to extracellular matrix accumulation and glomerular sclerosis over time (6, 15, 16). In addition, dysfunction of mesangial cells may further contribute to the reduced glomerular filtration observed in our model, as these cells play a crucial role in maintaining glomerular capillary structure and regulating the filtration surface area through their contractile properties (33). Collectively, these findings underscore the utility of this model for studying vascular-specific mechanisms of renal injury, independent of medullary developmental defects.

In *Ren1^ctdTomato^* Homo mice, vascular remodeling develops early with marked CAAH observed at two months of age despite preserved blood pressure. This dissociation between structural vascular changes and hemodynamic load supports a mechanism driven by cell-autonomous alterations in renin-lineage cells, rather than by pressure-induced stress. Although circulating Ang I and Ang II levels are already reduced at this early stage, residual Ang II likely sustains vascular tone through compensatory mechanisms (34, 35). Over time, however, this balance deteriorates: by six months, Ang I and Ang II levels decline further, coinciding with worsening vascular hypertrophy and remodeling. These progressive structural changes may impair vascular compliance and autoregulation, contributing to the delayed blood pressure reduction observed at later stages.

The concentric arteriolar hypertrophy phenotype observed in these mice is similar to the one observed in animals with deletion of renin globally or constitutively in renin cells (9, 13). However, the latter shows additional renal abnormalities, including papillary atrophy, underdeveloped medulla, hydronephrosis, interstitial fibrosis, focal glomerulosclerosis, and perivascular infiltration of mononuclear cells. These morphological alterations can be the result of developmental changes caused by the absence of renin expression throughout the life of the animals and may alter the vascular phenotype. Unlike mice with global deletion of renin, *Ren1^ctdTomato^* Homo mice do not require neonatal saline injections to survive and do not develop other kidney abnormalities, indicating that the bicistronic approach rendered a renin hypomorphic mouse.

In conclusion, *Ren1^ctdTomato^* Homo mice constitute an excellent model for the study of the mechanisms involved in the development of concentric vascular hypertrophy under RAS inhibition. In addition, this model may provide a better understanding of factors controlling renin protein folding, stability, packaging, and release.

## AKNOWLEDGEMENTS

We thank Xiuyin Liang, Fang Xu, Minghong Li, and DJ White for excellent technical assistance.

